# Derivation of predator functional responses using a mechanistic approach in a natural system

**DOI:** 10.1101/2020.11.04.368670

**Authors:** Andréanne Beardsell, Dominique Gravel, Dominique Berteaux, Gilles Gauthier, Jeanne Clermont, Vincent Careau, Nicolas Lecomte, Claire-Cécile Juhasz, Pascal Royer-Boutin, Joël Bêty

**Author notes:** Corresponding authors; Andréanne Beardsell and Joël Bêty.

## Abstract

The functional response is central to our understanding of any predator–prey system as it establishes the link between trophic levels. Most functional responses are evaluated using phenomenological models linking predator acquisition rate and prey density. However, our ability to measure functional responses using such an approach is often limited in natural systems and the use of inaccurate functions can profoundly affect the outcomes of population and community models. Here, we develop a mechanistic model based on extensive data to assess the functional response of a generalist predator, the arctic fox (*Vulpes lagopus*), to various tundra prey species (lemmings and the nests of geese, passerines and sandpipers). We found that predator acquisition rates derived from the mechanistic model were consistent with field observations. Although sigmoidal functional responses were previously used to model fox-prey population dynamics, none of our simulations resulted in a saturating response in all prey species. Our results highlight the importance of predator searching components in predator-prey interactions, especially predator speed, while predator acquisition rates were not limited by handling processes. By combining theory with field observations, our study provides evidences that predator acquisition rate is not systematically limited at the highest prey densities observed in a natural system. We reinforce the idea that functional response categories, typically types I, II, and III, should be considered as particular cases along a continuum. Specific functions derived with a mechanistic approach for a range of densities observed in natural communities should improve our ability to model and understand predator-prey systems.

## Introduction

A long-standing problem in ecology is to measure how the consumption rate of a predator varies with prey availability. The functional response is at the core of predator-prey theory as it establishes the link between predator acquisition rate and prey density (Solomon 1949). Functional response shapes are typically categorised as linear (type I), hyperbolic (type II) or sigmoidal (type III; Holling 1959a,b). This classification is commonly used by ecologists when incorporating predation into population and community models (Fryxell et al. 2007; Serrouya et al. 2015; Turchin and Hanski 1997), and type II is the most widely applied model (Rall et al. 2012). The shape of the functional response can have major consequences on the outcomes of population and community models. For instance, a type III promotes stability or coexistence whereas a type II destabilizes predator-prey dynamics (Murdoch 1973; Sinclair et al. 1998). Describing the functional response of pairwise trophic interactions is also important to understand higher-order interactions. For instance, the shape of the functional response alone can profoundly change predictions about the outcome of predator-mediated trophic interactions (Abrams et al. 1998; Holt and Bonsall 2017).

Quantifying and determining the shape of functional responses remains an important challenge, especially in natural systems. Most empirical research on functional responses has been conducted under controlled laboratory or field enclosure conditions (96%, *n* = 116 studies, reviewed by Pawar et al. 2012) where prey density is manipulated, predator consumption is recorded, and the functional response models are compared through statistical analysis. In natural systems, our ability to measure functional response is limited by a combination of factors: small sample size, a relatively narrow gradient of observed prey densities, the difficulty to observe predator-prey interactions directly, or the difficulty to estimate predator and prey numbers (Ellis et al. 2019; Gilg et al. 2006; Suryawanshi et al. 2017; Therrien et al. 2014). The large variability around predator acquisition rates can also constrains our ability to fully discriminate among functional response shapes, and hence limits our ability to accurately model predator-prey interactions in complex and natural ecosystems (Chan et al. 2017; O’Donoghue et al. 1998; Vucetich et al. 2002). Moreover, phenomenological models fail to identify the proximate mechanisms regulating predator acquisition rates.

Derivation of functional responses based on measurable features of species behaviour (e.g. speed, attack and success probability) provides several advantages. Compared with phenomenological models, mechanistic models 1) allow assessing the shape of the functional response based on behavioral attributes of the predator, 2) are based on parameters with a direct biological interpretation, and hence have the potential to reinforce links between theory and data (Connolly et al. 2017). The number of mechanistic models of predator-prey interactions is growing, and most of them aim to predict trophic links based on species traits, especially body size (Gravel et al. 2013; Ho et al. 2019; Portalier et al. 2019). Mechanistic models of functional response further allow the integration of predator-prey pairs to describe trophic links, which can improve our ability to model complex ecological interactions.

The main objective of our study is to develop a mechanistic model to characterize and quantify functional responses of a generalist mammalian predator to various prey species. The originality of our approach is to assess functional response i) by focusing on four components of predation (searching, chasing, capturing, and handling prey) and ii) by using field experiments and detailed behavioral observations to parameterize each step of the mechanistic model. We evaluated the coherence of our models using data from a long-term field study that estimated prey densities and predator acquisition rates. We also performed sensitivity analyses to identify the main proximate drivers of change in predator acquisition rates. Finally, we modeled the potential effects of density dependence in components of predation on the shape of the functional responses within the range of prey densities observed in the field.

The mechanistic model was developed for the arctic fox (*Vulpes lagopus*), a generalist predator of the tundra ecosystem, using highly detailed empirical observations from a long-term ecological monitoring program in the Arctic (Gauthier et al. 2013). This system offers several benefits to study predator-prey interactions among vertebrates, including a relatively simple food web, an open landscape and the continuous summer daylight allowing direct behavioral observations. The arctic fox is an active hunting predator that travels extensive daily distances within its territory in summer (M.-P. Poulin and D. Berteaux, unpublished manuscript). Lemmings and birds (mostly eggs and juveniles) are the main components of the summer diet of arctic foxes in most tundra ecosystems (Angerbjörn et al. 1999; Giroux et al. 2012). Lemmings exhibit population cycles with peak density every 3-5 years (Fauteux et al. 2015), and the arctic fox predation pressure on tundra ground-nesting birds is typically released at high lemming density (Bêty et al. 2002; McKinnon et al. 2014; Summers et al. 1998). Surprisingly, the exact mechanisms driving this well-known short-term apparent mutualism between lemmings and birds are still unclear, but they likely involve fox functional responses (Bêty et al. 2002; Flemming et al. 2016; Summers et al. 1998).

A few studies attempted to quantify the functional responses of arctic fox using phenomenological models (Angerbjörn et al. 1999; Eide et al. 2005; Gilg et al. 2006). Relatively low sample sizes reduced the ability of previous studies to fully distinguish between different shapes of functional responses. Moreover, the hoarding behavior of arctic foxes was not considered in previous estimations of functional responses (Angerbjörn et al. 1999; Eide et al. 2005; Gilg et al. 2006). Like many other animals (Vander Wall 1990), arctic foxes can predate more prey than they consume on the short-term, and such behavior can strongly increase prey acquisition rates, e.g. foxes foraging in goose colonies can hoard between 40% and 97% of eggs acquired during the bird nesting period (Careau et al. 2008; Samelius and Alisauskas 2000). Although type III functional responses were previously used to model fox-prey population dynamics (Gilg et al. 2003, 2009), food hoarding may substantially reduce handling time and could therefore make the shape of the functional response linear or slightly convex (Oksanen et al. 1985).

## Methods

### Study system

During the summer, the southwest plain of Bylot Island, Nunavut, Canada (73° N; 80° W) harbors a large greater snow goose colony (*Anser caerulescens atlanticus*; ~20,000 pairs). Insectivorous migratory birds are also nesting in the study area and include the lapland longspur (*Calcarius lapponicus*), a passerine, and several species of shorebirds (primarily *Calidris* spp. and *Pluvialis* spp.). Two species of small mammals are present, the brown (*Lemmus trimucronatus*) and collared (*Dicrostonyx groenlandicus*) lemmings. The brown lemming has high-amplitude cycles of abundance with a 3–5-year periodicity, whereas the collared has low-amplitude cycles (Gruyer et al. 2008). The mammalian predator guild is dominated by the arctic fox and the ermine (*Mustela erminea*). The arctic fox is the main nest predator of geese (Bêty et al. 2002; Lecomte et al. 2008), sandpipers (McKinnon and Bêty 2009; Royer-Boutin 2015) and passerines (Royer-Boutin 2015). Additional details on plant communities and general landscape can be found in Gauthier et al. (2013).

The model was parametrized and evaluated using data from Bylot Island, where foxes and their prey have been monitored since 1993. We observed foraging foxes using binoculars and spotting scopes (20 x 60x) from one or two blinds located in the middle of the goose colony during 10 summers between 1996 and 2019.

### Mechanistic model of functional responses

We used the Holling disk equation as a starting point to build the mechanistic model of functional response (Holling 1959a) inspired by the general formalism of Pawar et al. (2012). Predation was broken down into four different processes, which are searching, chasing, capturing, and handling of a prey item by a predator. Acquisition rate of a prey item (species *i*) by a predator (*f* (*i*)), namely the functional response, takes the following form:

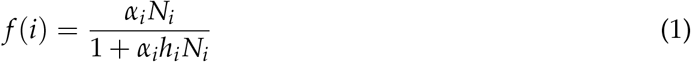

where *α_i_* is the capture efficiency (km^2^/h), *N_i_* the prey density (number of *i*/km^2^), and *h_i_* the handling time of prey (h/*i*). Capture efficiency is obtained by the product of predator speed (*s*; km/h), reaction distance (*d_i_*; km), detection (*z_i_*) and attack probability (*k_i_*) of the prey by the predator, and the success probability (*p_i_*) of an attack (table 1):

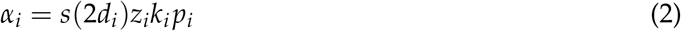

**Table 1:**
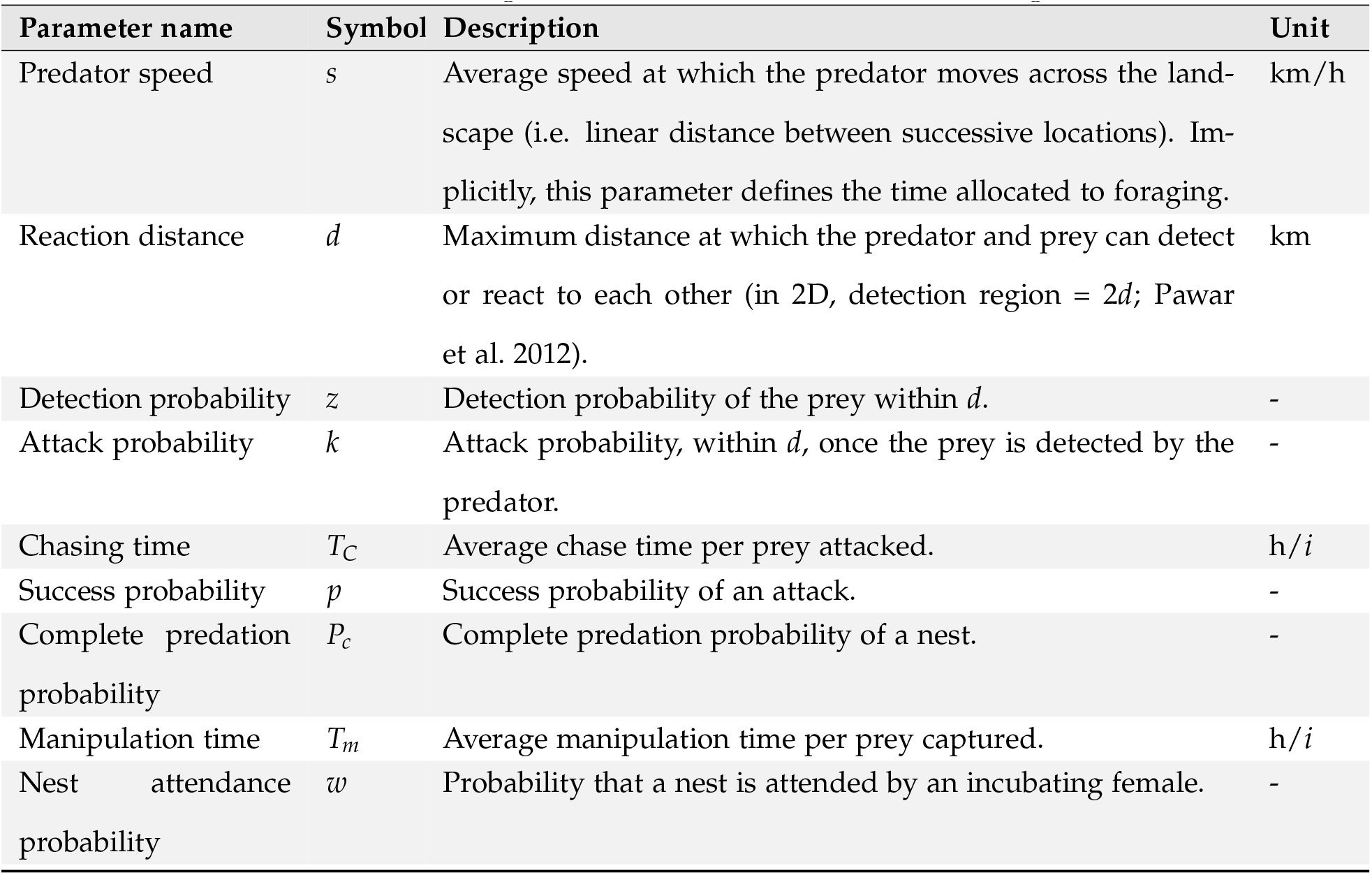
Definition of the parameters used in the functional response model.

The combination of the time spent chasing the prey once encountered 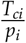 and the time spent manipulating the prey once subdued *T_mi_* define an overall prey handling time (*h_i_*):

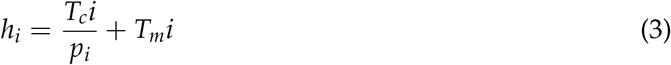

*α_i_* depends only on prey density, and we assumed that prey are randomly distributed. Satiety was not considered as a potential mechanism limiting acquisition rate. Indeed, foxes can predate more prey than they consume on the short-term; e.g. about 4% (*n* = 128) and 48% (*n* = 98) of predated eggs and lemmings are immediately eaten, respectively (Careau et al. 2007). Predator interference was not incorporated in the model as foxes rarely encounter and interact with other individuals while foraging within their summer territory (49 interactions, which represents 0.9% of the time over 118 hours of direct observations of foxes foraging in the study area). The full model derivation is provided in Appendix C.

The general model of functional response (eq. 1) allows for a continuum between a linear and a hyperbolic functional response shape. In order to allow the model to extend to a sigmoidal shape, we added density dependence in capture efficiency components that were expected to vary with prey density (i.e. reaction distance and detection and attack probabilities; see below).

### Prey specific functional responses

We adapted the general model (eq. 1) to each prey species based on their life history traits and anti-predator behavior (fig. 1). The specific models for each prey species are provided in table C2.

**Figure 1:**
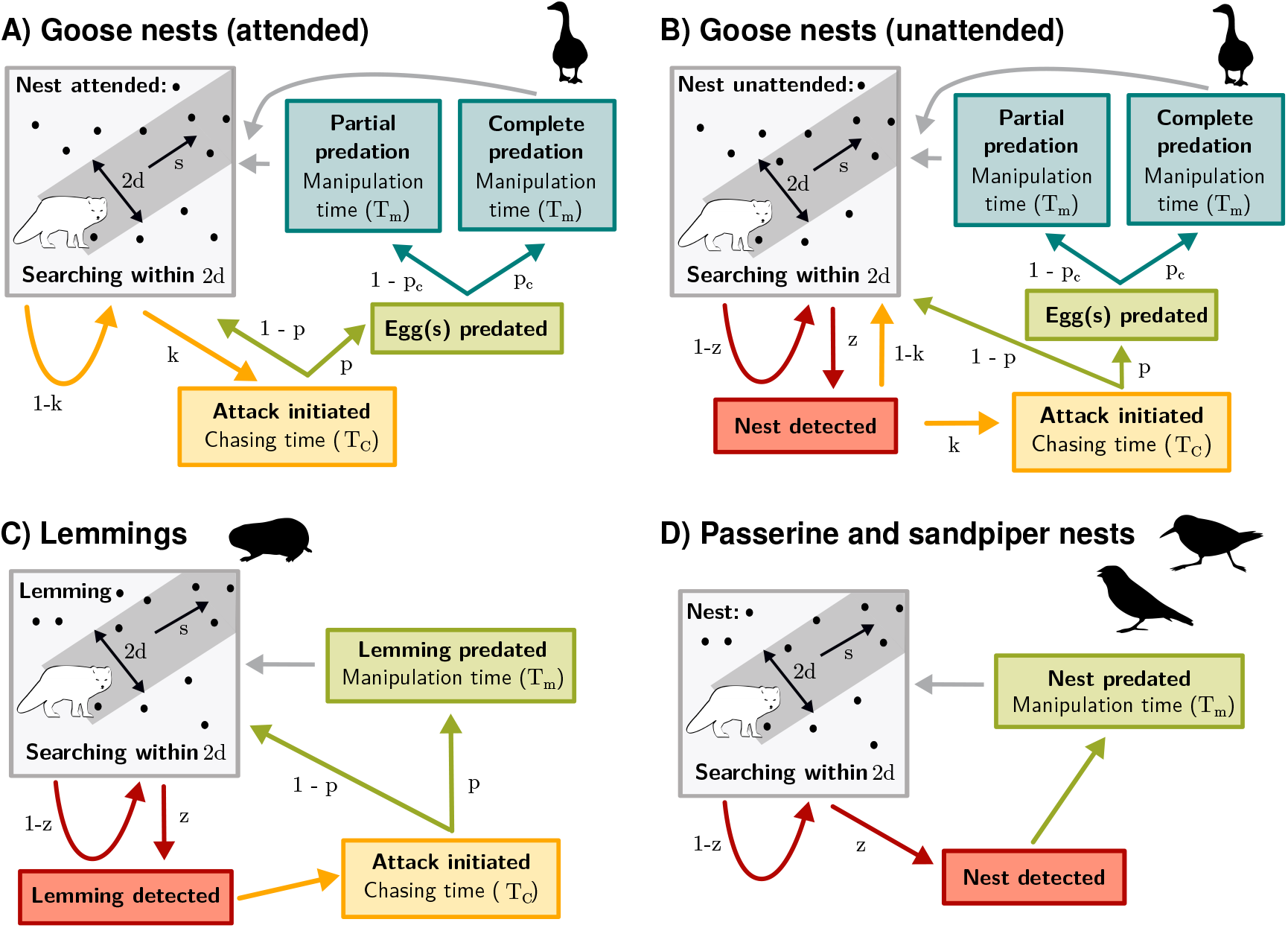
Conceptual mechanistic model of functional response of arctic fox to each prey species: attended (**A**) or unattended (**B**) goose nests, lemmings (**C**), and passerine and sandpiper nests (**D**). Parameters are as follows : *d* is the reaction distance, *s* the predator speed, *z* the detection probability, *k* the attack probability, *p* the success probability, *P_c_* the probability of complete nest predation, *T_C_* the time spent chasing, and *T_m_* the time spent manipulating the prey.

For goose nests, the first modification to the general model was to add a component for complete and partial nest predation. This modification was necessary since a successful attack by the predator does not always result in complete clutch predation (Bêty et al. 2002), which can affect manipulation time and, ultimately, acquisition rates. The second modification to the general model was to split the model into two components. A first component models acquisition rate of goose nests when the female is incubating or when one protecting adult is at < 10 m from the nest (attended nest; fig. 1A). A second component models acquisition rate of goose nests during incubation recesses when both adults are at >10 m from the nest (unattended nest; fig. 1B). Geese can actively protect their nests against arctic foxes; therefore, their presence at the nest strongly influences fox foraging behavior (Bêty et al. 2002; Samelius and Alisauskas 2001), which translates into changes in capture efficiency components. Parameter values of capture rates were thus estimated separately for goose nests that were attended or unattended by an incubating female (table C1). When a nest is attended by a highly conspicuous snow goose, we assumed that nest detection probability is 1 within *d* (fig. 1A). For unattended nests, we used a detection probability function obtained from an artificial nests experiment (fig. C1). Sometimes, unattended nests can be protected if parents detect a fox during an incubation recess and return quickly to their nest. Like attended nests, we thus estimated success probability (*p*) and complete clutch predation probability (*P_c_*) for unattended nests (fig. 1B). The third and last modification was to introduce the nest attendance probability (*w*). We estimated this parameter by combining information on the average time spent on the nest by females and on the average distance between females and their nest during the goose incubation period (Poussart et al. 2000; Reed et al. 1995; see Appendix C).

The general model (eq. 1) was simplified for lemmings as we assumed that an attack is systematically initiated by the fox once a lemming is detected within *d* (fig. 1C). The general model was also simplified for passerines and sandpipers as we assumed that once a passerine or a sandpiper nest is detected, the nest is always predated (fig. 1D).

We incorporated density dependence into the goose and the lemming models within the range of densities observed in our study system. For each parameter in which density dependence was incorporated, the minimum and the maximum parameter values were associated respectively with the minimum and the maximum prey density in order to calculate the slope and the intercept of the density-dependence relationship. In the goose model, we modified attack and success probabilities for attended nests, and reaction distance and detection probability for unattended nests. In the lemming model, we added density dependence in reaction distance and detection probability. The rationale behind these additions is that predators may form search images for abundant prey, which can increase their ability to detect them (Ishii and Shimada 2010; Nams 1997). As predators could also increase their attack rate and success as prey density increases, we added density dependence in attack and success probabilities. We did not incorporate density dependence into the passerine and sandpiper nest models as the range of nest densities observed in our study system is likely too low to influence fox behavior (maximum of 12 nests/km^2^ compared to a maximum of 926 goose nests and 414 lemmings per km^2^). See Appendix C for more details on the incorporation of density dependence.

The model was implemented in R v. 3.6.0 R Core Team 2019.

### Parameter values

The model was parameterized mostly using data from Bylot Island but also from the literature when data were missing. Parameters were derived from field experiments using artificial nests or estimated using arctic fox GPS tracking data and direct observations of foraging foxes (table C1). See Appendix C for a detailed description of the method used to extract each parameter.

### Evaluating the coherence between the mechanistic model and empirical predator acquisition rates

Predator acquisition rates at different prey densities were assessed in the field annually using two independent methods. These data did not allow validation of the shape of the functional responses, but they provided a way to evaluate the performance of the mechanistic model in estimating prey acquisition rates at the various prey densities observed in our study system.

First, we obtained goose eggs and lemming acquisition rates by conducting direct observations of foraging foxes for 10 summers between 1996 and 2019 during the goose incubation period (details on behavorial observations can be found in Bêty et al. (2002) and Careau et al. (2008)). For each year, the acquisition rate was calculated as the total number of prey acquired (goose eggs or lemmings) divided by the total length of the observation bouts of individual foxes. The acquisition rate of a clutch of eggs was estimated by dividing the acquisition rate of goose eggs by the annual average clutch size. For the years where information was available, we also calculated the acquisition rate for attended and unattended nests. We estimated annual goose nest density either by visual counts of the nests located in the observation zone (range: 0.5-3 km^2^) during the incubation period (1996-1999, 2019) or over a fixed 0.2 km^2^ plot within the intensively monitored core area of the goose colony (2004-2005, 2015-2016). We estimated lemming density annually with snap traps from 1994 to 2009 and with live traps from 2004 to 2019 (see Fauteux et al. 2018 for methods). We summed the density estimate of brown and collared lemming.

Second, we obtained passerine and sandpiper nest acquisition rates by monitoring annually (2005 to 2013) the fate of passerine and sandpiper nests (Gauthier et al. 2013; McKinnon et al. 2014). Nest density was estimated as the number of passerine and sandpiper nests found in a 8 km^2^ plot systematically searched throughout the breeding season. We estimated acquisition rate of nest content (eggs or chicks) by using the daily survival rate of nests (*dsr*), the total number of nests found in the study plot (*N_tot_*), the number of foxes foraging in the plot (*N_fox_*) and the proportion of nests predated by foxes (*P_fox_*). Since foxes establish territorial pairs on Bylot (Rioux et al. 2017), we assumed that 2 foxes were foraging in the study plot. We also considered that foxes were responsible for 100% (*n* = 19) and 81% (*n* = 25) of the failed sandpiper and passerine nests, respectively, as indicated by camera monitoring (McKinnon and Bêty 2009; Royer-Boutin 2015). An estimation of the acquisition rate is obtained by:

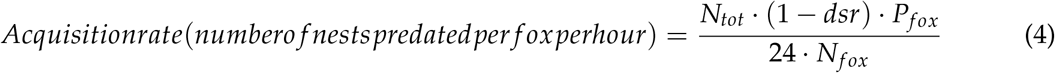

The daily nest survival rate was modeled using the logistic exposure method (Shaffer 2004). Additional details on daily nest survival rate calculations and nest monitoring methods can be found in Royer-Boutin (2015). Density estimates for all prey species were standardized as the number of nests per km^2^.

### Uncertainty and sensitivity analysis

We quantified how uncertainty in parameter values affected estimation of predator acquisition rates by using the Latin hypercube sampling technique (an efficient implementation of the Monte Carlo methods; Marino et al. 2008). This analysis allowed us to investigate the uncertainty in the model output generated by the uncertainty in parameter inputs. Each parameter was represented by a probability distribution (uniform or normal truncated) based on the distribution of empirical data (table C1). For some parameters, the biological information was limited, so we assigned a uniform distribution allowing for a large range bounded by minimum and maximum values. Latin hypercube sampling was then applied to each distribution (*N* = 1000 iterations). This method involved dividing a probability distribution into *N* equal probability intervals that were then sampled without replacement, resulting in *N* iterations of the model using each combination of parameters values. This method allowed us to explore the entire range of each parameter and most of them encompass various environmental conditions (e.g. weather conditions, prey availability). We computed the median, the 90, the 95, and 99 percentiles of the model output by using the empirical cumulative distribution.

We also conducted a local sensitivity analysis to identify key parameters of the mechanistic models within the range of prey densities observed in our study system. We modified each parameter value by ± 100% while holding others constant, and we assessed how this variation affected the predator acquisition rate (expressed as % of change).

## Results

From 1996 to 2019, we observed foraging foxes in the goose colony for 124 hours. Average goose nest density was 409 nests/km^2^ (range: 100-926 nests/km^2^) and lemming density was 193 ind./km^2^ (range: 11-414 ind./km^2^; table B1). Average acquisition rates were 0.61 nest/fox/h for goose nests and 0.94 ind./fox/h for lemmings. The majority of eggs acquired (67%) were from unattended nests, while 33% were from attended nests (*n* = 218). Average passerine nest density was 7.7 nests/km^2^ (range: 6.1-12.3 nests/km^2^), and sandpiper density was 2.5 nests/km^2^ (range: 1.0-5.9 nests/km^2^; table B2). Average acquisition rates were 0.10 nest/fox/h and 0.04 nest/fox/h for passerine and sandpiper nests, respectively.

The uncertainty analysis revealed that varying all parameters simultaneously generated considerable variation in model output (fig. 2). Nonetheless, no parameter combinations resulted in a saturating functional response for all prey species within the range of prey densities observed in our study system: the acquisition rate at maximal prey density was below the saturation point in all simulations (see histograms in fig. 2 and fig. A1). Based on the value of the parameters estimated within the observed prey densities, acquisition rate at saturation was 8 nests/fox/h for goose nests, 17 ind./fox/h for lemmings, 166 nests/fox/h for passerine and 26 nests/fox/h sandpiper nests. About 90% (fig. 2A), 100% (fig. 2B and 2D), and 89% (fig. 2C) of the empirical estimations fall within the 99 percentile of the predator acquisition rates derived from the mechanistic models. As the goose nest model was split for attended and unattended goose nests, we also computed acquisition rates separately for each of these situations. Goose nest acquisition rate was higher for unattended nests than attended nests, which is consistent with empirical estimations (fig. A2). Although most (66%) of the empirical estimations fell within the 95 percentiles of the model, all field estimates for unattended nests were under the model median at nest densities above 200 nests/km^2^ (fig. A2). This may indicate a slight overestimation of the proportion of unattended nests at relatively high densities.

**Figure 2:**
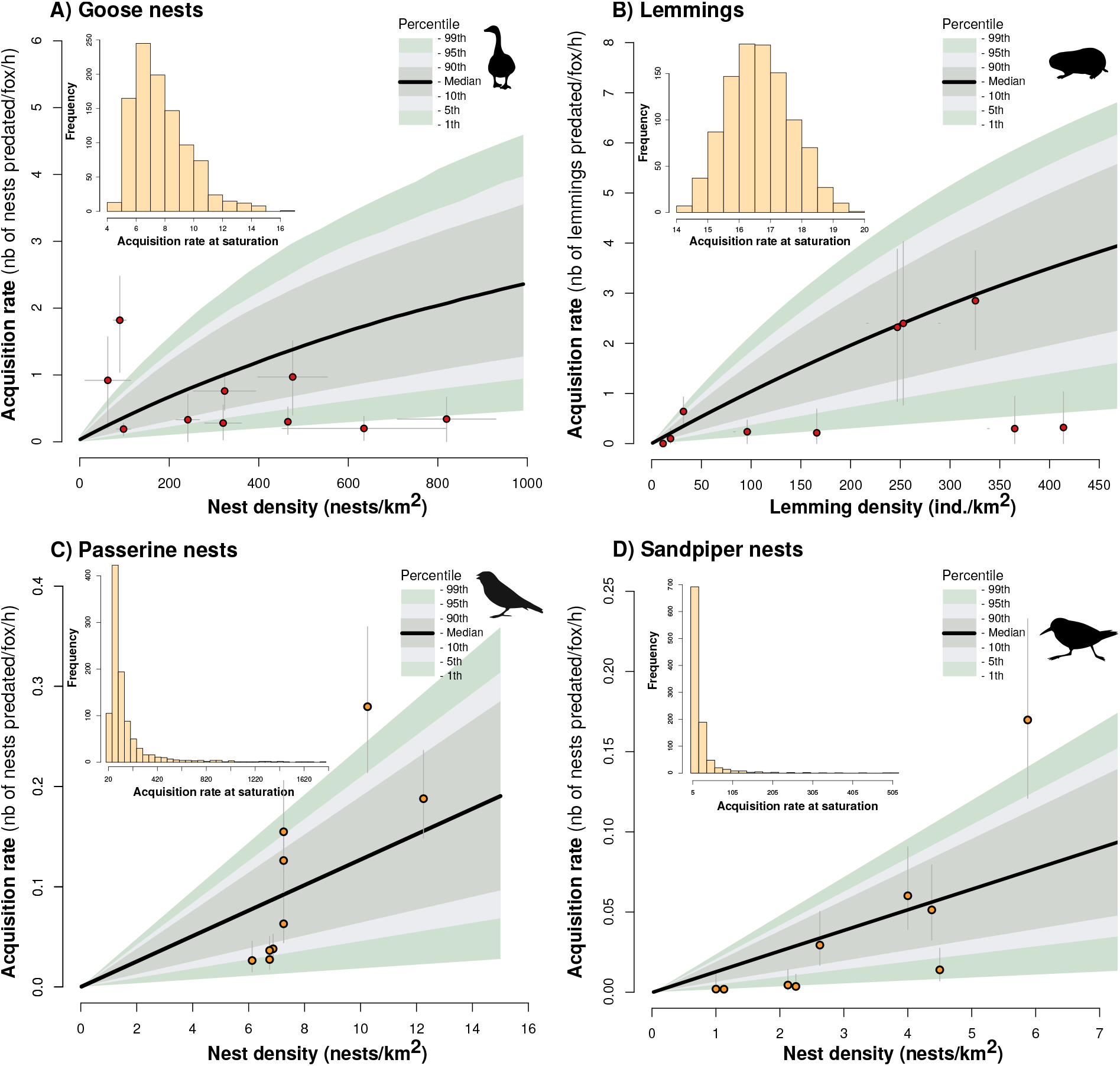
Functional response of arctic fox to density of goose nests (**A**), lemmings (**B**), passerine nests (**C**), and sandpiper nests (**D**). Black lines represent the median of the mechanistic model and the color bands represent the 90, 95, and 99 percentiles based on 1000 simulations. Empirical data are represented by red and yellow dots respectively. Histograms in the inset show the distributions of acquisition rate at saturation for each simulation. Horizontal error bars in (**A**) indicate the range of nest density during the incubation period. Vertical errors bars in (**A**) and (**B**) represent standard errors calculated using bootstrapping. Errors bars in (**C**) and (**D**) represent 95% confidence intervals from daily survival rate estimates.

Predator speed was an influential parameter of the functional response of all prey species (fig. 3). Goose nest acquisition rate was generally more affected by parameters associated with unattended nests than attended nests (fig. 3A). The magnitude of change in goose nest acquisition rate related to the changes in manipulation time increased slightly with nest density. Lemming acquisition rate was not affected by chasing and manipulation time, whereas detection distance, and detection and success probability had an influence equivalent to predator speed (fig. 3B). Similarly, functional response models of passerine and sandpiper nests were not sensitive to change in manipulation time, whereas detection distance and detection probability had an influence equivalent to predator speed (fig. 3C).

**Figure 3:**
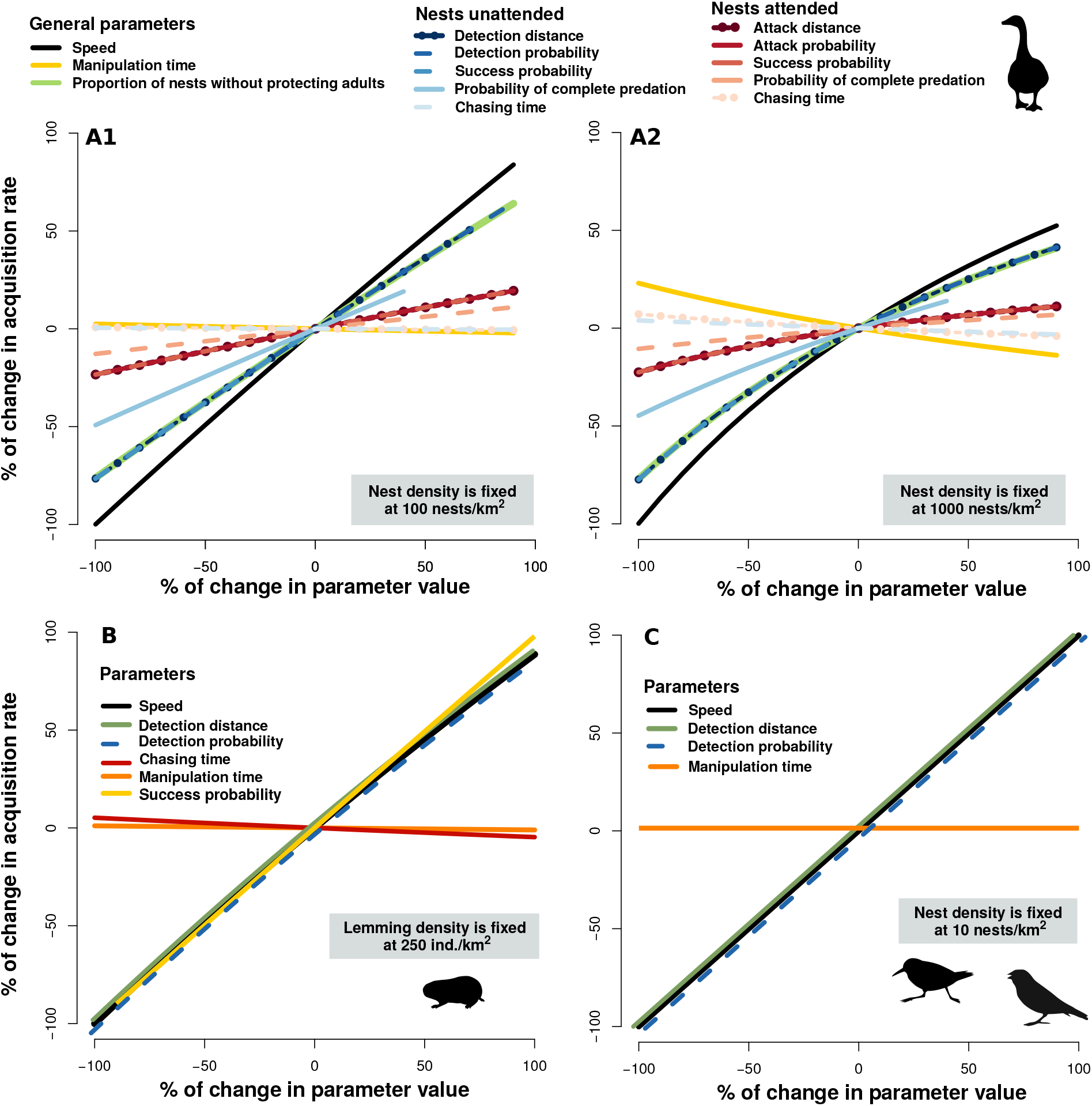
Sensitivity of predator acquisition rates to changes in parameter values of the mechanistic models used to assess the functional response of arctic fox to goose nests (at 100 (**A1**), and 1000 nests/km^2^ (**A2**)), to lemmings at 250 ind./km^2^ (**B**) and to passerine and sandpiper nests at 10 nests/km^2^ (**C**).

Even though the shape of the functional response changed slightly between models without or with density dependence in capture efficiency components (allowing for a gradient between type I and type III), a maximum difference of 1.4 nests/fox/h at 1000 goose nests/km^2^ and 1.3 lemmings/fox/h at 450 lemmings/km^2^ were found between models (fig. 4).

**Figure 4:**
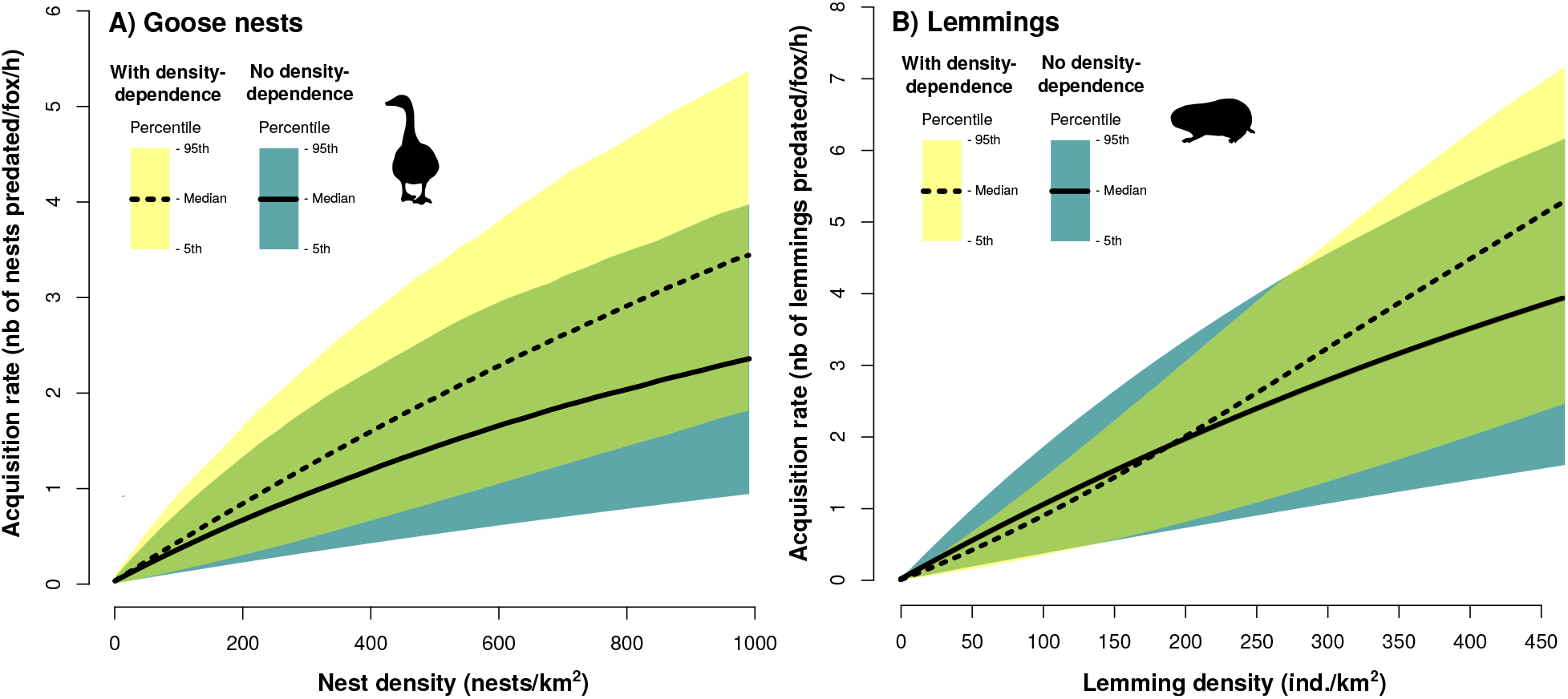
Functional response models of arctic fox to goose nests (**A**) and lemmings (**B**) with and without density dependence on capture efficiency components, within the range of densities observed in the field. Black lines represent the median of the mechanistic model and the color bands represent the 95 percentiles based on 1000 simulations.

## Discussion

Most models of predator-prey interactions make assumptions regarding the shape of the functional response expressed by predators. By quantifying the characteristics of a predator (e.g. speed and detection distance), we developed a mechanistic model of arctic fox functional response to various prey species. Our model derives the shape of the functional response along a gradient from linear to sigmoidal. We benefited from extensive empirical data on all prey species to parameterize and evaluate the coherence of the model. Predator acquisition rates derived from the mechanistic model were consistent with field observations, and the main proximate mechanisms driving predator acquisition rates were also identified. In all prey species, predator speed was an influential parameter, while handling time was not a limiting process. Although type III functional responses were previously used to model fox-prey population dynamics (Gilg et al. 2003, 2009), our simulations indicate that predator acquisition rate was not systematically limited at the highest prey densities observed in our study system. Our model allows for a mechanistic interpretation of the functional response of predator-prey pair and could be extended to more complex modules involving multiple predators and prey species.

Holling’s functional response models (type II and III), which are commonly used in population dynamics models (Gervasi et al. 2012; Serrouya et al. 2015; Turchin and Hanski 1997), predict that predator acquisition rates should eventually saturate at high prey densities. In our study system, we found no evidence of arctic fox saturation at the highest prey densities observed. Several factors may explain this result. First, the hoarding behaviour of arctic fox may substantially reduce handling time by limiting the constraints associated with digestion and satiety, which can make the functional response shape linear or slightly convex even at high prey densities (Oksanen et al. 1985). Second, while predator acquisition rates must theoretically become constrained by handling and/or digestion at high prey densities, the prey densities required to reach a saturation point could be rarely observed in natural systems. Indeed, empirical support for saturating functional response in the wild is relatively rare and comes mostly from controlled laboratory experiments in which the range of prey densities may exceed the range observed in nature (99% of all type II functional response were derived from controlled laboratory experiments (*n* = 61 studies); reviewed by Rall et al. 2012). Such an issue can be avoided when mechanistic approaches are used to derive functional responses. One particularity of our system is the presence of a large goose colony where prey density can be quite high (up to ~900 nests/km^2^). Interestingly, even in this context, we found no evidence of predator saturation. Hence, our results add to a growing body of research indicating that predators may not become systematically satiated at the highest densities of prey observed in nature (Chan et al. 2017; Novak 2010; Preston et al. 2018).

Historically, a categorical approach was adopted by ecologists to define functional responses. A linear functional response was traditionally attributed to filter feeders (Jeschke et al. 2004), a hyperbolic shape (type II) to invertebrates and a sigmoidal shape (type III) to vertebrate predators (Holling 1965). Although, this categorization has some heuristic value in introductory texts and can be useful in some aspects of research where categorization is necessary, types I, II, and III should be considered simply as particular cases along a continuum. Instead of using a priori shapes to describe functional responses, our study illustrates how mechanistic models can generate functions linking prey density and predator acquisition rates that are specific, and hence more relevant, to the range of densities observed in a food web. Considering the strong effect of functional responses on the outcome of predator-prey models (Abrams et al. 1998; Sinclair et al. 1998), such specific functions should improve our ability to adequately simulate and quantify the strength of direct and indirect species interactions in natural communities.

We did not incorporate predator dependence in the functional response model, despite a growing body of studies indicating that some mechanisms (e.g. facilitation, interference) are likely to occur in functional responses (Novak et al. 2017). However, arctic foxes maintained summer territories (averaging 9.6 km^2^) with low overlap (A. Grenier-Potvin and D. Berteaux, unpublished manuscript), which prevents potential interference within territories. We are thus confident that variation in predator density should not affect our main conclusions. Nonetheless, the mechanistic model could be extended to more complex predator-prey systems, including predator interference.

Habitat characteristics could affect several parameters of the mechanistic model, hence the functional response shape and magnitude could be modulated by the structural complexity of the landscape (Barrios-O’Neill et al. 2015; Toscano and Griffen 2013). For instance, the detection distance of a nest by arctic foxes could be lower in dense vegetation (Flemming et al. 2016), the attack probability could be lower for nests located in wetlands and islets only accessible by swimming (Gauthier et al. 2015; Lecomte et al. 2008), and the success probability of an attack could be modulated by the presence of complex networks of lemming tunnels offering refuges. Exploration of the effects of structural complexity on functional responses remains rare (but see Barrios-O’Neill et al. 2015; Lipcius and Hines 1986; Toscano and Griffen 2013), and more empirical research is needed to integrate these sources of variation in mechanistic models.

The outputs of the mechanistic model were generally consistent with field observations. However, adding more complexity could improve its performance and our ability to identify the main drivers of predator acquisition rates. For instance, group defence and mutual vigilance are additional factors that may reduce predator acquisition rates at high prey density (Clark and Robertson 1979). Although there is no evidence of group defence in geese (Bêty et al. 2001), the snow goose could benefit from the vigilance and early warning provided by neighbors nesting nearby (Samelius and Alisauskas 2001). Beyond a threshold of goose nest density, such anti-predator behavior could reduce the proportion of unattended nests with increasing nest densities. Nest attendance probability was an influential parameter of the goose model and mutual vigilance may partly explain why acquisition rates observed in the field at moderate-high nest densities were under the model median (fig. 2A).

One mechanism often advanced for explaining the apparent mutualism between two prey sharing a common predator is predator saturation or satiation (Abrams and Matsuda 1996; Holt 1977). Our results showed that the arctic fox is not satiated at the highest lemming densities observed in our study system. This suggests that the underlying mechanism for the short-term positive effect of high lemming density on arctic bird reproductive success (Bêty et al. 2002; Blomqvist et al. 2002) is likely not predator satiation nor saturation, and it reinforces the need to adopt a mechanistic approach to fully understand predator-prey interactions. Short-term positive indirect effect could arise from changes in various components of the functional response. For instance, attack probability of an attended goose nest could be inversely dependent of lemming density, or daily distance traveled by the predator (speed) could vary according to prey availability. As indicated by our sensitivity analyses, attack probability was not a strong driver of prey acquisition rates while predator speed was an important parameter affecting all prey species. Hence, lemming-induced changes in predator speed due to changes in reproductive state and activity budget of foxes could be an alternative hypothesis explaining the apparent mutualism between lemmings and arctic birds. To fully identify the main proximate drivers of indirect interactions in natural communities, we need to identify the components of capture rate for a given prey that change according to variation in densities of other prey and evaluate the impact of such changes on prey mortality using mechanistic models.

## Conclusion

Previous studies of functional responses typically tried to discriminate between predetermined shapes of functional responses. Our study illustrates how mechanistic models based on empirical estimates of the main components of predation can generate functional responses specific to a range of prey densities relevant to a given food web. Such mechanistically derived functional responses are needed to untangle proximate drivers of predator-prey population dynamics and to improve our understanding of predator-mediated interactions in natural communities. Although it would be unrealistic to resolve every pairwise interaction within ecological networks, our mechanistic model provides a starting point for studying higher-order effects such as indirect interactions that can emerge among prey species.

## Acknowledgments

The research relied on the logistic assistance of the Polar Continental Shelf Program (Natural Resources Canada) and of Sirmilik National Park of Canada. The research was funded by (alphabetical order): Arctic Goose Joint Venture, the Canada Foundation for Innovation, the Canada Research Chairs Program, the Canadian Wildlife Service, the Fonds de recherche du Québec-Nature et technologies, the International Polar Year program of Indian and Northern Affairs Canada, the Natural Sciences and Engineering Research Council of Canada, the ArcticNet Network of Centers of Excellence, the Northern Ecosystem Initiative Program (Environment Canada), the Northern Scientific Training Program, the Nunavut Wildlife Management Board, Polar Knowledge Canada, Université du Québec à Rimouski, Université Laval and the W. Garfield Weston Foundation. We are especially grateful to the many people who helped us with field work over many years, the Mittimatalik Hunters and Trappers Organization and Park Canada’s staff for their assistance.

## Appendix A: Supplementary Figures

**Figure A1:**
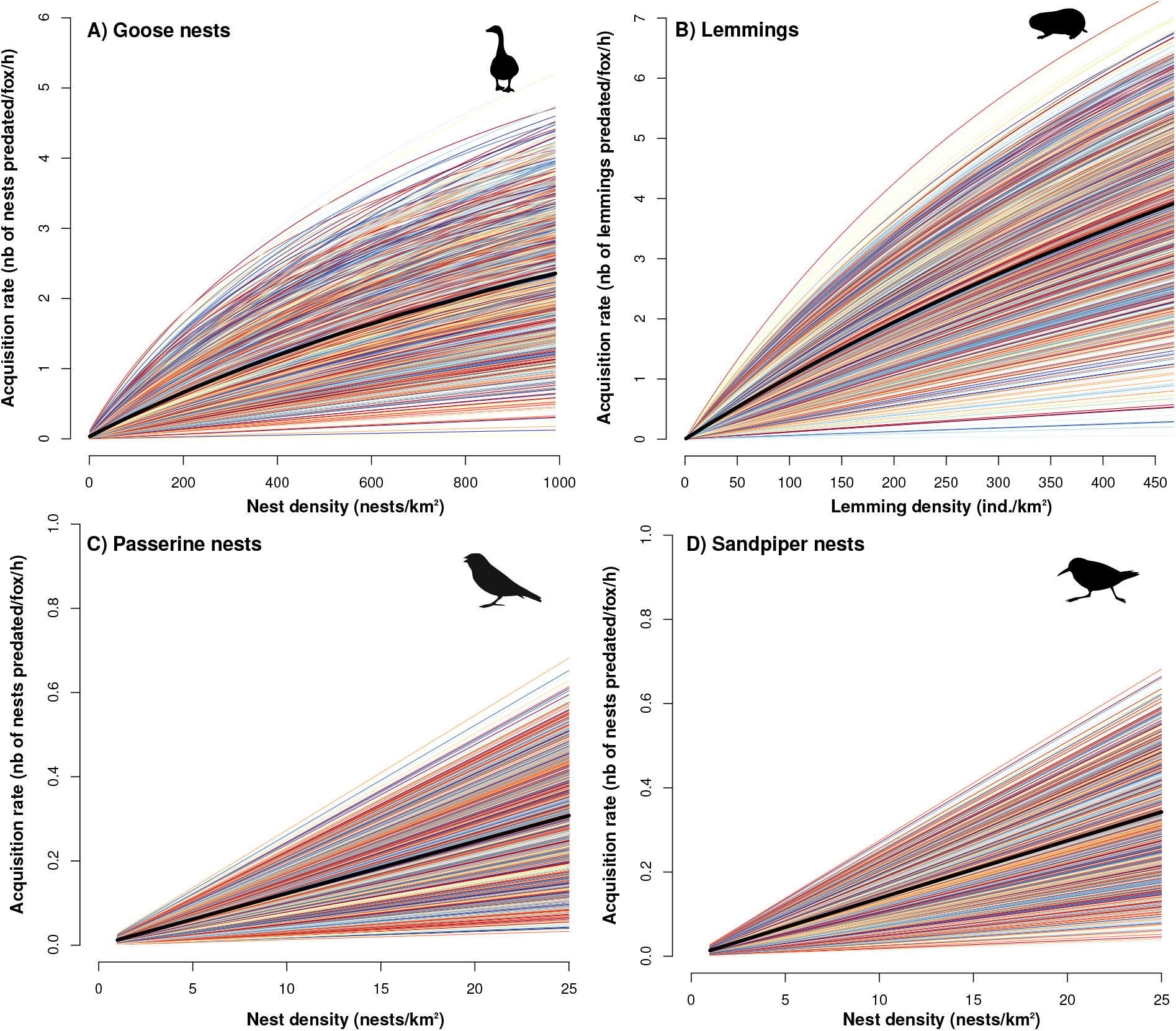
Functional response of arctic fox to density of goose nests (**A**), lemmings (**B**), passerine nests (**C**), and sandpiper nests (**D**). Each line reprents a simulation and the solid black line represents the model median (*N* = 1000 simulations).

**Figure A2:**
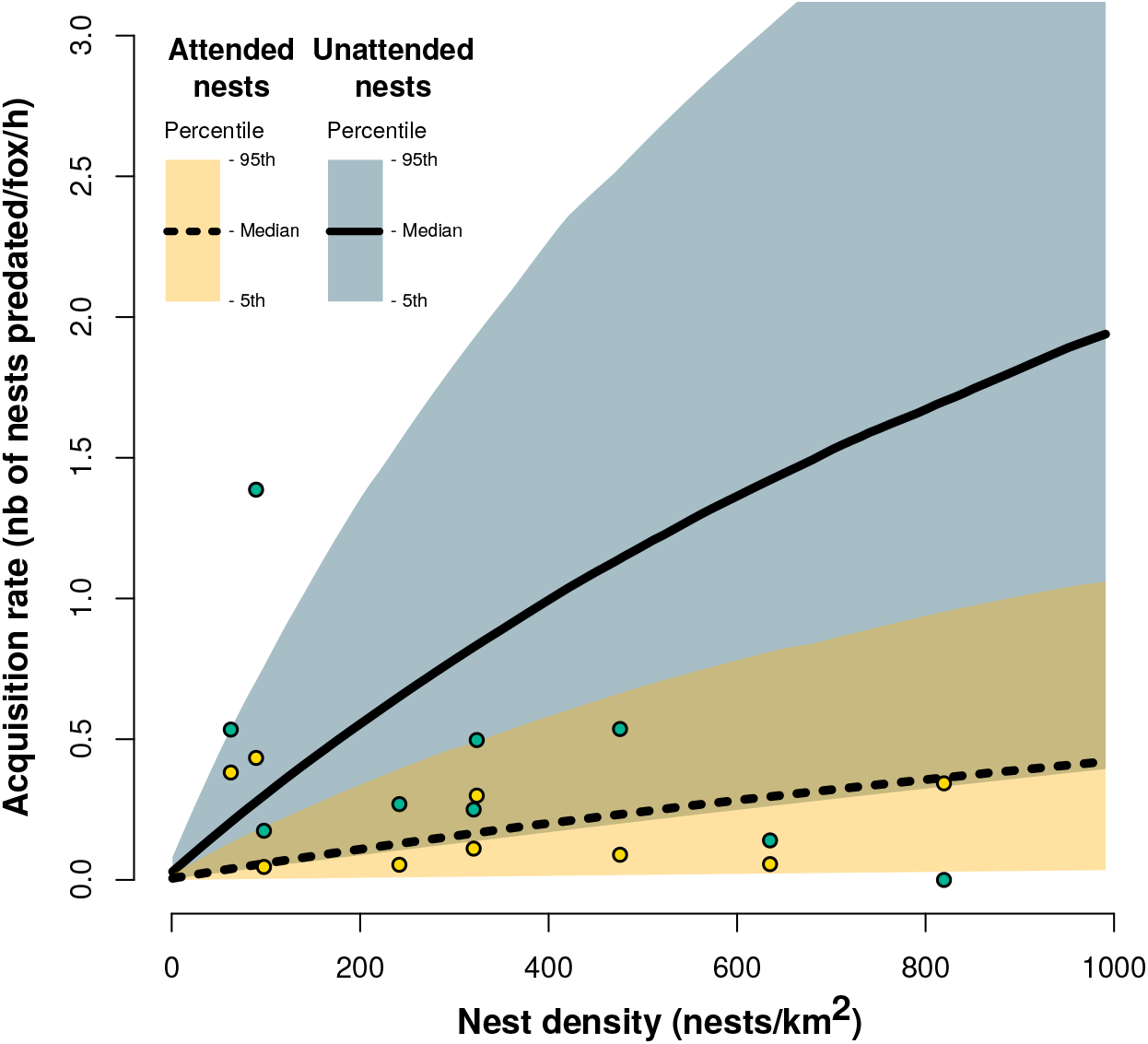
Functional response of arctic fox to density of goose nests for attended and unattended nests. The black lines represent the model median and the color bands represent the 95 percentiles based on 1000 simulations. Empirical data for attended and unattended nests obtained from direct observations of foraging foxes are represented by yellow and blue dots respectively.

## Appendix B: Supplementary tables

**Table B1:** Empirical data used to evaluate the performance of the mechanistic model of functional response of arctic fox to goose nests and lemmings on Bylot Island, Nunavut. Data are listed by year.

| Year | Observation length (h) | Average clutch size | Goose density range (min-max nests/km<sup>2</sup>) | Goose nest acquisition rate (nests/fox/h) | Lemming density (ind./km<sup>2</sup>) | Lemming acquisition rate (ind./fox/h) |
| --- | --- | --- | --- | --- | --- | --- |
| 1996 | 3.1 | 3.7 | 79-100 | 1.82 | 414 | 0.32 |
| 1997 | 4.7 | 3.9 | 219-264 | 0.33 | 166 | 0.21 |
| 1998 | 9.5 | 3.8 | 283-358 | 0.28 | 247 | 2.32 |
| 1999 | 4.2 | 3.1 | 15-110 | 0.92 | 11 | 0 |
| 2004 | 29.8 | 3.6 | 85-110 | 0.19 | 325 | 2.85 |
| 2005 | 43.6 | 3.6 | 257-390 | 0.76 | 32 | 0.64 |
| 2015 | 3.4 | 3.5 | 713-926 | 0.34 | 365 | 0.30 |
| 2016 | 3.3 | 3.4 | 401-550 | 0.97 | 253 | 2.40 |
| 2017 | 10.1 | 3.5 | 456-814 | 0.20 | 19 | 0.10 |
| 2019 | 12.7 | 4.0 | 381-465 | 0.30 | 137 | 0.24 |

**Table B2:** Empirical data used to evaluate the performance of the mechanistic model of functional response of arctic fox to passerine and sandpiper nests on Bylot Island, Nunavut. Data are listed by year.

| **Year** | **Passerine**<br>**nest density**<br>**(nests/km<sup>2</sup>)** | **Passerine nest**<br>**acquisition rate**<br>**(nests/fox/h)** | **Sandpiper**<br>**nest density**<br>**(nests/km<sup>2</sup>)** | **Sandpiper**<br>**acquisition rate**<br>**(nests/fox/h)** |
| --- | --- | --- | --- | --- |
| 2005 | 7.3 | 0.15 | 4.4 | 0.051 |
| 2006 | 10.3 | 0.28 | 5.9 | 0.169 |
| 2007 | 6.1 | 0.03 | 2.1 | 0.004 |
| 2008 | 7.3 | 0.06 | 1.0 | 0.002 |
| 2009 | 6.9 | 0.04 | 1.1 | 0.002 |
| 2010 | 6.8 | 0.04 | 4.5 | 0.014 |
| 2011 | 6.8 | 0.03 | 2.3 | 0.004 |
| 2012 | 12.3 | 0.19 | 4.0 | 0.060 |
| 2013 | 7.3 | 0.13 | 2.6 | 0.029 |

## Appendix C: Additional Methods

### Derivation of the mechanistic model of functional response

The area searched (*A*; km^2^) by a predator is expressed by the product of predator speed (*s*, km/h), the reaction distance to a prey item *i* (*d_i_*, km), and the time spend searching (*T_s_*, h):

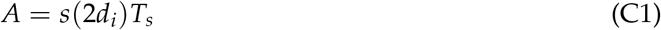

A potential encounter occurs when the predator comes within the distance (*d_i_*) at which one can detect and react to the other. As not all prey within this area may be detected, attacked and subdued by the predator, we introduced the detection probability (*z_i_*), the attack probability (*k_i_*), and the success probability of an attack (*p_i_*). Capture efficiency of a prey item *i* by the predator is expressed by:

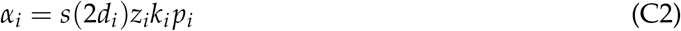

The number of prey captured (*V_αi_*) during a search duration *T_S_* for a density *N_i_* is:

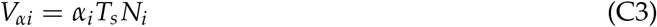

The time spent searching (*T_S_*) is defined as:

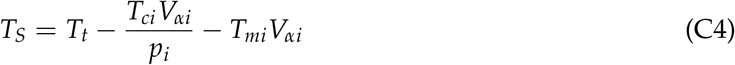

where *T_t_* is the time available for feeding; 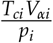 is the time spent chasing prey once they are encountered; and *T_mi_V_αi_* is the time spent manipulating prey if they are subdued.

By simplifying Eq. C4 we have:

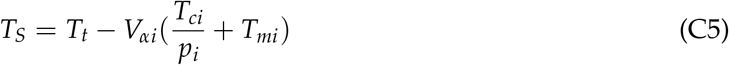

We can combine the chasing and manipulation time to produce an overall prey handling time (*h_i_*):

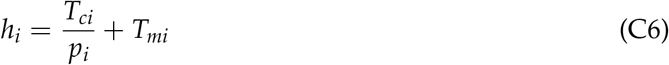

Substituting *T_s_* from Eq. C5 into Eq. C3, we arrive at:

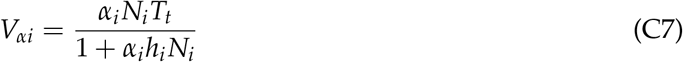

The functional response of a predator (*f* (*i*)) is the number of prey captured per predator per unit of time. This is expressed by dividing Eq. C7 by *T_t_*:

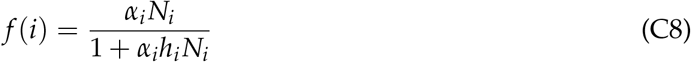

### Estimation of parameter values

#### Predator speed

A total of 16 foxes (7 females and 9 males) were equipped with a GPS collar (Radio Tag-14, Milsar, Poland) during the summers of 2018 and 2019. Of those, 7 were equipped during both years giving us a total of 23 fox-summers (8 foxes in 2018 and 15 in 2019). Reproductive females (2 out of 4 in 2018 and 6 out of 7 in 2019) spent substantial time nursing the pups in the den. Foxes were captured using Tomahawk cage traps #205 (Tomahawk Live Trap Company, Tomakawk, WI, USA) or Softcatch #1 padded leghold traps (Oneida Victor Inc. Ltd., Cleveland, OH, USA) as described in (Rioux et al., 2017). GPS fix intervals were fixed at 4 minutes (360 fixes per day), and data were downloaded to a hand-held receiver using UHF transmission. GPS location error was of 11 m (M.-P. Poulin and D. Berteaux, unpublished manuscript).

Average predator speed (km/day) was estimated by adding linear distances between successive locations, using the *adehabitatLT* library in R (R Core Team, 2019). Predator speed was extracted from June 5 to July 9 to focus on the incubation period of most birds. We removed from analyses all fixes obtained <48 hours after capture and days where the number of fixes was insufficient (<75% of all fixes). Predator speed was converted per hour and average speed was 1.52 km/h (sd = 0.59 km/h, *n* = 123 fox-days).

#### Goose parameters

##### Nest attendance probability

During goose incubation period, the time spent on the nest by females average 93% (*μ* = 93.6, se = 1.6%, *n* = 7 females; Poussart et al. 2000 and *μ* = 93, *n* = 41 females; Reed et al. 1995). During incubation recesses females usually remained close to their nests, and 90% of all records (*n* = 183) were within 20 m (Reed et al., 1995). Since there is uncertainty in the proportion of females within 10 m, we used 90% as the maximum and 50% as the minimum. By combining this information, we can estimate a minimum probability of nest attendance at 96.5.% and a maximum probability of nest attendance at 99.3%.

##### Detection probability

We used artificial nests to assess experimentally the detection probability of unattended goose nests in summer 2019. Goose eggs were simulated with domestic hen eggs. Two eggs were placed in each artificial nest and covered with goose down collected from old goose nests. A total of 24 paired artificial nests were deployed randomly in the goose colony. The two paired nests were separated either by 10, 30, 60, 80 or 100 m. Movement-triggered cameras (model PM35T25, Reconyx) were set directly on the ground 5 m from each nest allowing to identify nest predators and to determine the exact time of nest predation. A successful detection was considered to have occurred when the two paired nests were predated by a fox in the same time interval (*n* = 9 cases). An unsuccessful detection occurred when only one nest of the pair was depredated by a fox (*n* = 6 cases). When a nest was predated by another species (e.g. gulls, ravens) or the event of predation was undetected by the camera, the pair was excluded from the analysis (*n* = 9 cases). We used a linear model with a binomial distribution to model detection probability in relation to detection distance (fig. C1).

**Figure C1:**
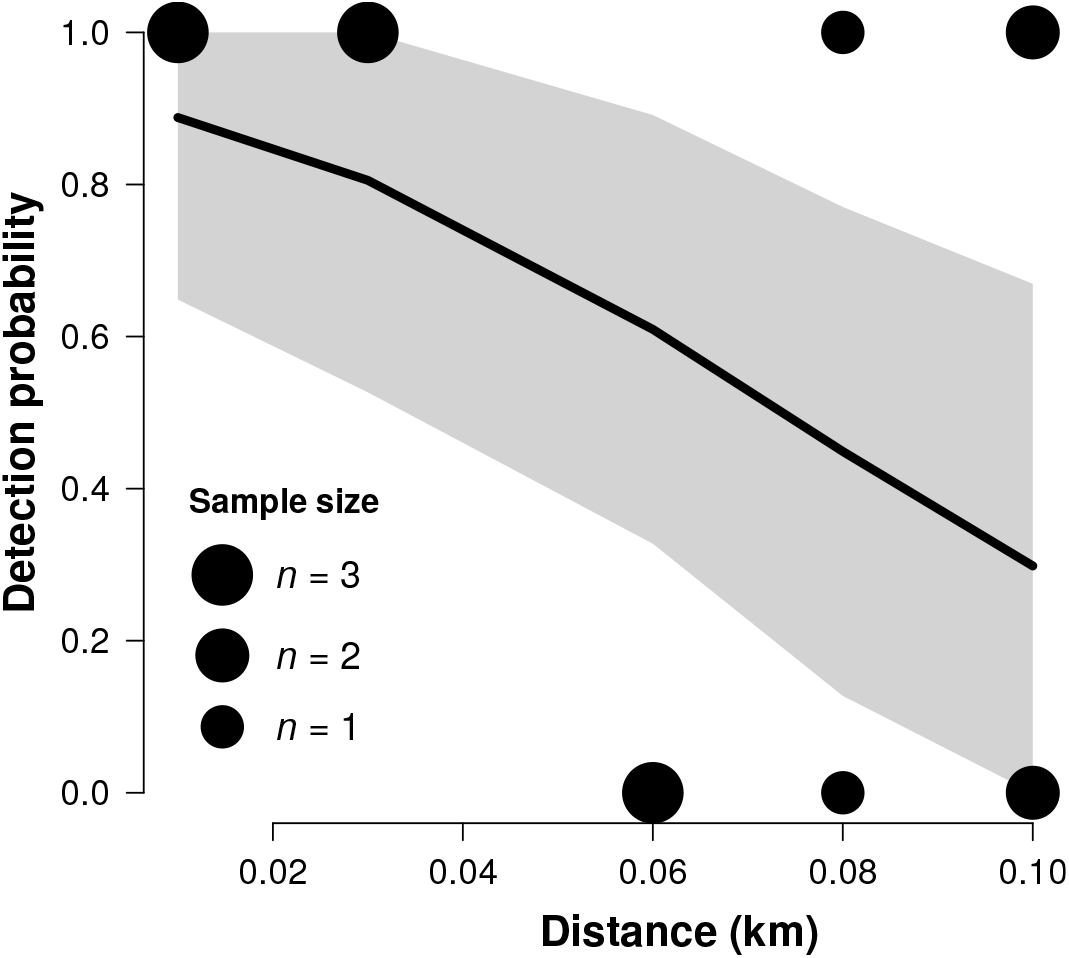
Probability that an artificial (unattended) goose nest was detected and depredated by an arctic fox as a function of distance. Each circle represents observed values and circle size is proportional to the number of observations. The curve represents the average detection probability and the gray band represents the 95% confidence interval of the regression.

##### Reaction distance

The reaction distance on attended nests was defined as the distance at which an attack can be initiated by the predator. This parameter was estimated with direct observations of foraging foxes in summer 2019 (*μ* = 0.0328, se = 0.007, *n* = 25 attacks). On unattended nests, the reaction distance was defined as the maximum distance at which the predator can detect an unattended nest. We used a combination of direct observations combined with artificial nests (same experience as for the detection probability) to estimate the reaction distance on unattended nests (*μ* = 0.0365 km, se = 0.009, max = 0.1 km, *n* = 13). We used 0.1 to 0.12 km as a range of maximum distance as the detection probability was still around 30% at 0.1 km (fig. C1). As our sample size was limited for attended and unattended nests, we assigned a uniform distribution for both parameters.

##### Chasing time

By conducting direct observations, we obtained the average chase time per egg attacked from a nest (*μ* = 23 sec/egg, sd = 29, *n* = 148 attacks). To convert per nest, we subsampled the data set and summed up 4 individual values. This total was then adjusted to the average clutch size of 3.7 (*μ* = 0.02, se = 0.0024 h per nest).

##### Manipulation time

By conducting direct observations, we obtained the time required by the predator to manipulate one egg, which includes consumption and hoarding time (*μ* = 136 sec/egg, sd = 127, *n* = 207). To convert per nest, we subsampled the data set and summed up 4 individual values. This total was then adjusted to the average clutch size of 3.7 (*μ* = 0.14, se = 0.009 h per nest).

##### Attack probability of attended nests

Based on direct observations of foraging foxes, we estimated attack probability at 22% by using the proportion of attended nests attacked by a fox divided by the total number of possible attacks (*n* = 215). The total number of possible attacks is obtained by adding the number of attacks towards attended nests and the number of times the foxes are chased by the geese without having initiated an attack. The latter gives us an estimation of the number of opportunities to initiate an attack. This estimation of attack probability gives us the upper limit of attack probability as the total number of possible attacks is most likely underestimated. Thus, we used a wide range of values (from 0.01 to 0.22) and a uniform distribution for this parameter.

##### Success probability

Based on direct observations of foraging foxes, the probability of a successful attack was 93.4% on unattended nests (se = 0.022, *n* = 137 attacks) and 9.8% on attended nests (se = 0.011, *n* = 701 attacks).

##### Complete predation probability

Based on direct observations of foraging foxes, the probability of a complete predation once an egg is subdued on unattended nests was 69% (se = 0.12, *n* = 16) and was 47% (se = 0.13, *n* = 15) on attended nests.

#### Lemming parameters

##### Reaction distance and detection probability

Based on direct observations, foxes generally initiate their attacks on lemmings within 5 m (on 29 attacks recorded in 1996-1999 and 2019). We assumed that the maximum reaction distance was twice that distance (10 m) and that detection probability followed a decreasing sigmoid function. Hence, detection probability was considered fairly high within 5 m radius (100-80%) and declined sharply between 5 and 10 m.

##### Chasing time

By conducting direct observations of foraging foxes, we estimated the average chase time per lemming attacked (*μ* = 88 sec, se = 7.1, *n* = 246 attacks).

##### Success probability

By conducting direct observations of foraging foxes, we estimated the probability of a successful attack on a lemming at 51% (se = 0.03, *n*=268 attacks).

##### Manipulation time

Based on direct observations of foraging foxes, we estimated the average manipulation time per lemming captured, which includes consumption and hoarding time (*μ* = 37 sec, se = 3.0, *n* = 93).

#### Passerine parameters

##### Reaction distance

The reaction distance was defined as the maximum distance between an observer (a simulated predator) and the nest when the bird flushes the nest. To measure the reaction distance, a human observer approached nests from a random bearing at normal walking speed (~4 km/h). The observer made several approaches until the bird leaves the nest. At each approach the distance between the observer and the nest was noted. Flush distance was recorded for 45 different nests in summer 2019 and ranged from 0 to 20 m (*μ* = 2.8 m, sd = 3.6 m, *n* = 77) The distance was measured either by pacing or with a GPS unit.

##### Detection probability

Detection probability was estimated by following the same method as reaction distance. Besides recording flush distances, the distance between the observer and the nest was noted even if the bird didn’t flush. We used a linear model with a binomial distribution to model detection probability (0; the bird didn’t flush, 1; the bird flushed) in relation to the minimum distance between the observer and the nest (*n* = 167, fig. C2).

**Figure C2:**
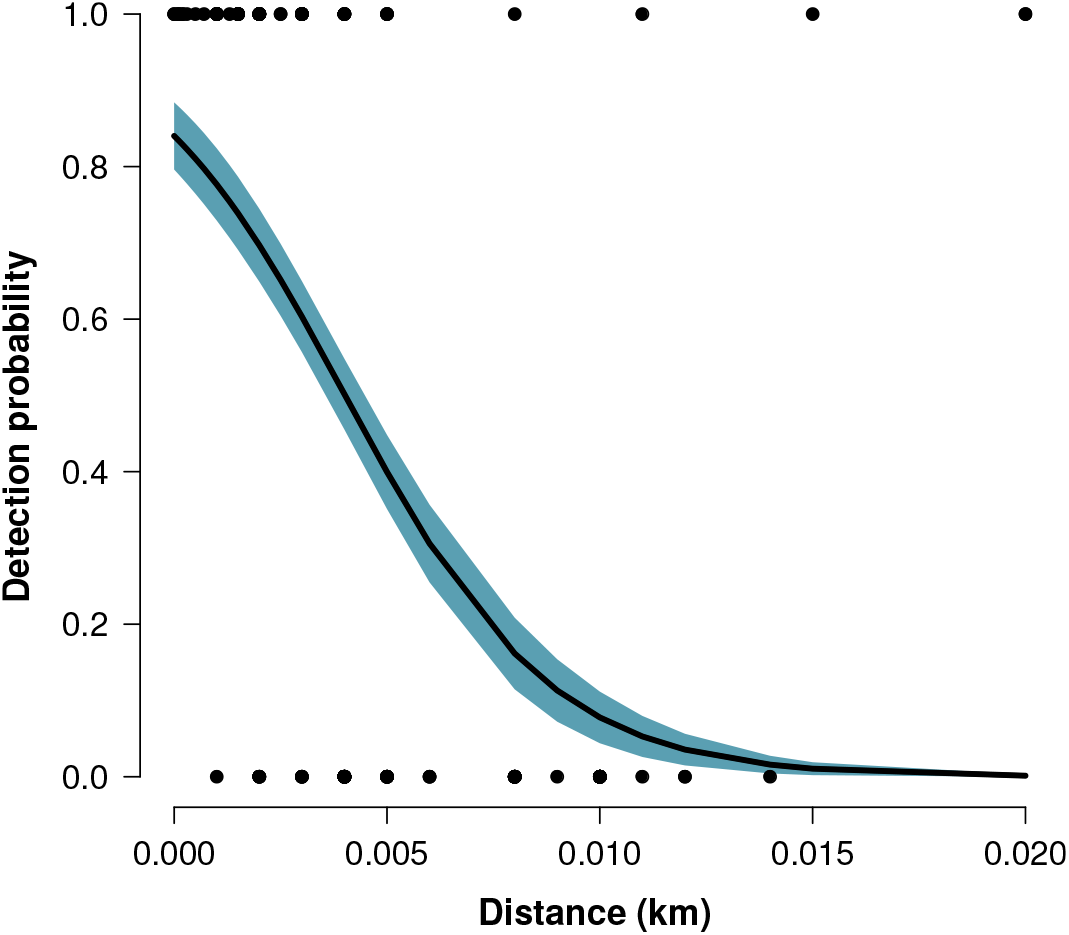
Detection probability of a passerine nest in relation to the distance between the observer and the nest. The gray band represents the ± standard error of the regression.

##### Manipulation time

By using movement-triggered cameras near the nest, we extracted manipulation time from 17 passerine nests monitored between 2006 and 2014. The average manipulation time was 31 sec (sd = 30 sec) and included consumption and hoarding time. Cameras (PM35T25 model) were set directly on the ground 5 m from the nest (for more details about camera features see McKinnon and Bêty (2009)).

#### Sandpiper parameters

##### Reaction distance and detection probability

Reaction distance was estimated by Smith and Edwards (2018) following the same method as described for passerines reaction distance. The mean reaction distance was 15.3 m (sd = 18.8 m, *n* = 104 nests, max = 85 m) for White-rumped Sandpiper (*Calidris fuscicollis*; Smith and Edwards (2018)). We used the same detection probability function as for passerines (fig. C2).

##### Manipulation time

By using movement-triggered cameras near real and artificial sandpiper nests, we extracted manipulation time from 5 sandpiper and 18 artificial nests monitored between 2006 and 2016. An artificial nest consisted of four Japanese quail (*Coturnix japonica*) eggs placed in a small depression made in the ground, similar to the simple nest scrapes used by sandpipers. Quail eggs resemble those of sandpipers in colouration and size. The average manipulation time was 250 sec (sd = 189 sec) and includes consumption and hoarding time. Cameras (PM35T25 model) were set directly on the ground 5 m from the nest (for more details about camera features see McKinnon and Bêty (2009)).

**Table C1:**
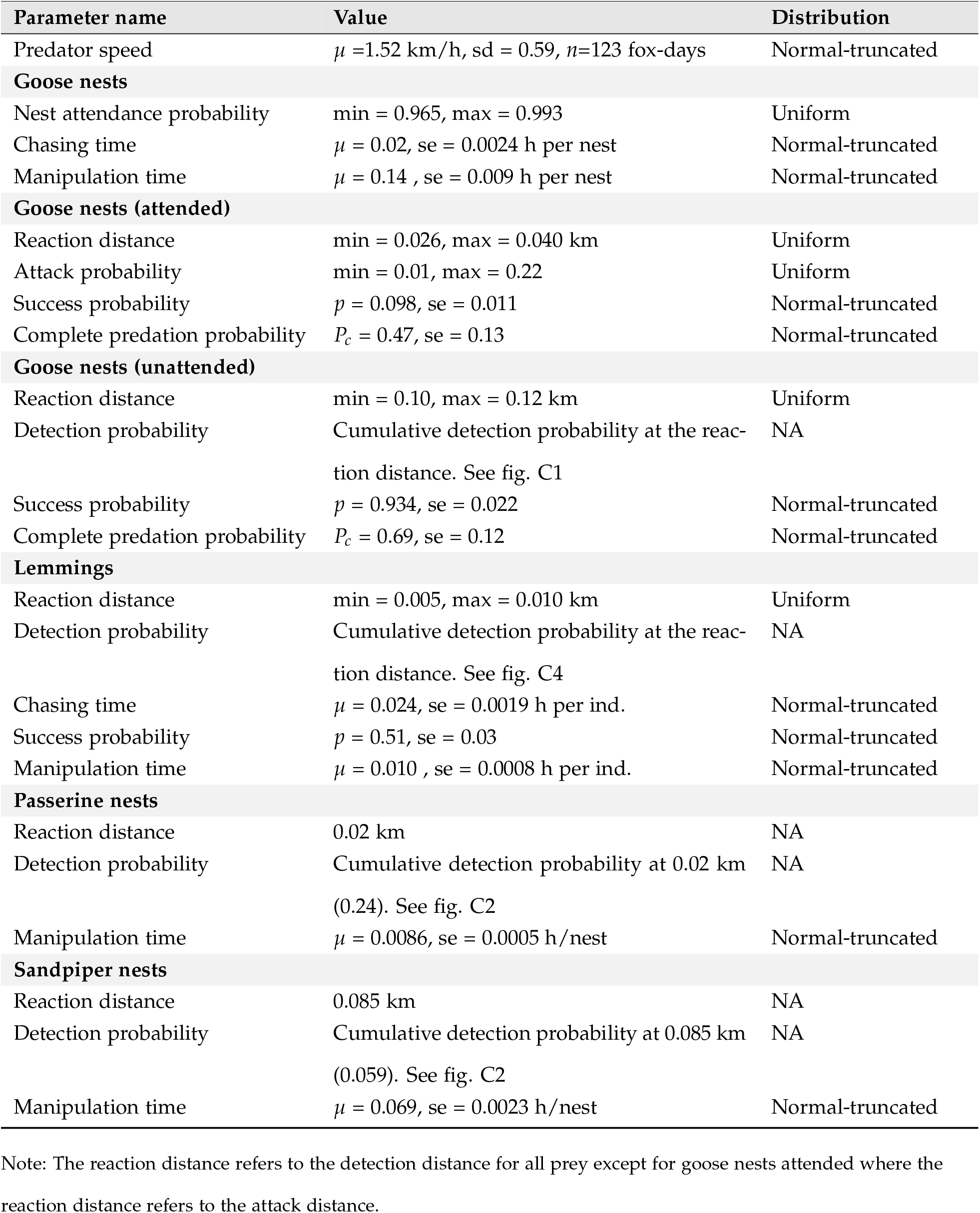
Parameter values and distribution used in the functional response model of arctic fox to goose nests, lemmings, passerine and sandpiper nests.

**Table C2:** List of equations to derive the functional response of arctic fox to each prey species.

| Model | Form |
| --- | --- |
| **Goose nests (G)** |  |
| Acquisition rate of G | $f(G) = f(G_a) + f(G_{ua})$ |
| **Goose nests (attended) ($G_a$)** |  |
| Capture efficiency of $G_a$ - Complete predation | $\alpha_{Gac} = s(2d_{Ga})k_{Ga}p_{Ga}P_{cGa}$ |
| Capture efficiency of $G_a$ - Partial predation | $\alpha_{Gap} = \frac{s(2d_{Ga})k_{Ga}p_{Ga}(1 - P_{cGa})}{3.7*}$ |
| Handling time of $G_a$ | $h_{Ga} = \frac{T_{cG}}{p_{Ga}P_{cGa}} + T_{mG}$ |
| Acquisition rate of $G_a$ | $f(G_a) = \frac{(\alpha_{Gac} + \alpha_{Gap})wN_G}{1 + (\alpha_{Gac} + \alpha_{Gap})h_{Ga}wN_G}$ |
| **Goose nests (unattended) ($G_{ua}$)** |  |
| Capture efficiency of $G_{ua}$ - Complete predation | $\alpha_{Guac} = s(2d_{Gua})z_{Gua}p_{Gua}P_{cGua}$ |
| Capture efficiency of $G_{ua}$ - Partial predation | $\alpha_{Guap} = \frac{s(2d_{Gua})z_{Gua}p_{Gua}(1 - P_{cGua})}{3.7*}$ |
| Handling time of $G_{ua}$ | $h_{Gua} = \frac{T_{cG}}{P_{cGua}} + T_{mG}$ |
| Acquisition rate of $G_{ua}$ | $f(G_{ua}) = \frac{(\alpha_{Guac} + \alpha_{Guap})(1 - w)N_G}{1 + (\alpha_{Guac} + \alpha_{Guap})h_{Gua}(1 - w)N_G}$ |
| **Lemmings (L)** |  |
| Capture efficiency of L | $\alpha_L = s(2d_L)z_Lp_L$ |
| Handling time of L | $h_L = \frac{T_{cL}}{p_L} + T_{mL}$ |
| Acquisition rate of L | $f(L) = \frac{\alpha_L N_L}{1 + \alpha_L h_L N_L}$ |
| **Passerine nests (P)** |  |
| Capture efficiency of P | $\alpha_P = s(2d_P)z_P$ |
| Handling time of P | $h_P = T_{mP}$ |
| Acquisition rate of P | $f(P) = \frac{\alpha_P N_P}{1 + \alpha_P h_P N_P}$ |
| **Sandpiper nests (S)** |  |
| Capture efficiency of S | $\alpha_S = s(2d_S)z_S$ |
| Handling time of S | $h_S = T_{mS}$ |
| Acquisition rate of S | $f(S) = \frac{\alpha_S N_S}{1 + \alpha_S h_S N_S}$ |

Note: \* Average clutch size of greater snow goose (Gauthier et al., 2013)

### Exploration of density dependence in capture efficiency components

We incorporated density dependence into the goose and the lemming models within the range of densities observed in our study system. For each parameter in which density dependence was incorporated, the minimum (*p_min_*) and the maximum (*p_max_*) parameter values were associated respectively with the minimum (*N_min_*) and the maximum (*N_max_*) prey density in order to calculate the slope and the intercept of the density-dependence relationship:

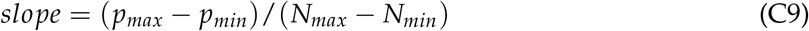

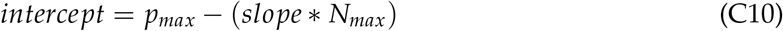

In the goose model, we modified attack and success probabilities for nests attended, and reaction distance and detection probability for nests unattended (e.g. Fig. C3). In the lemming model, we added density dependence in reaction distance, detection, attack and success probabilities (e.g. Fig. C4).

**Figure C3:**
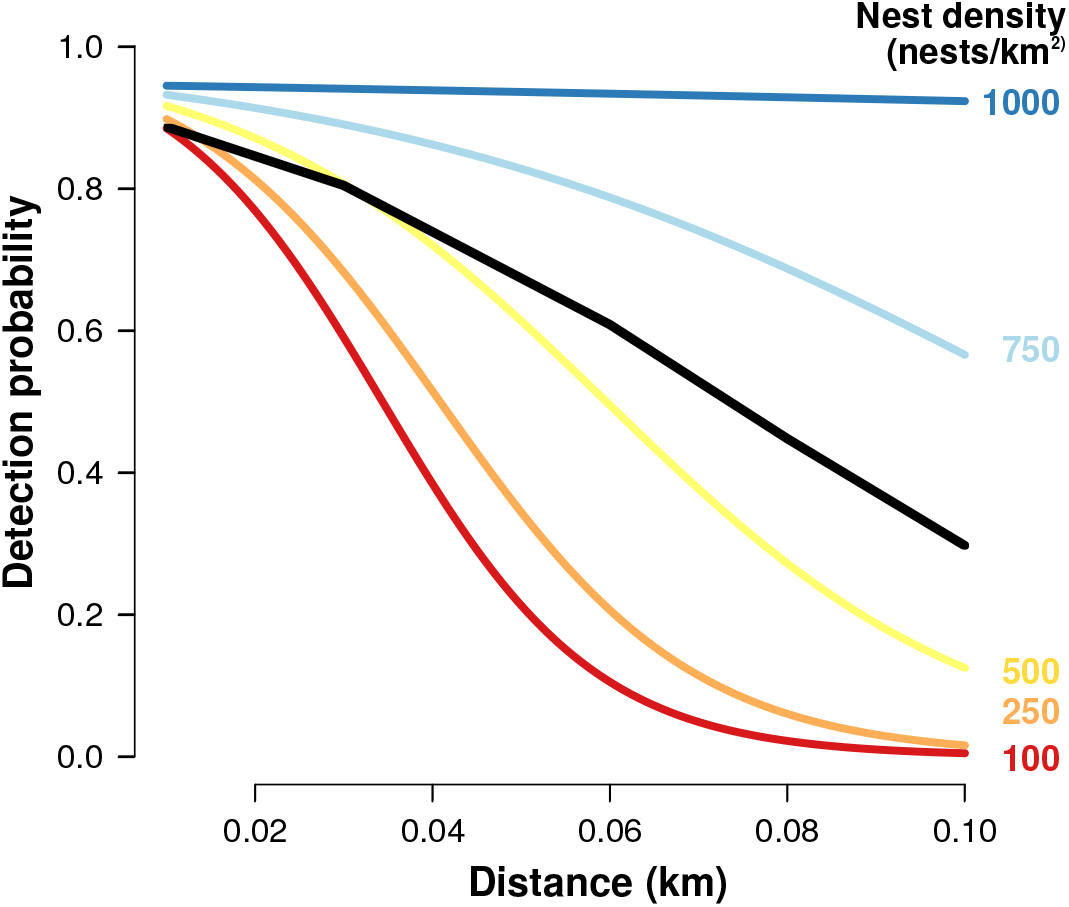
Probability that an artificial (unattended) goose nest was detected and depredated by the arctic fox as a function of distance along a gradient of goose nest density. The color gradient indicates the range of detection functions used to explore density dependence in detection probability.

**Figure C4:**
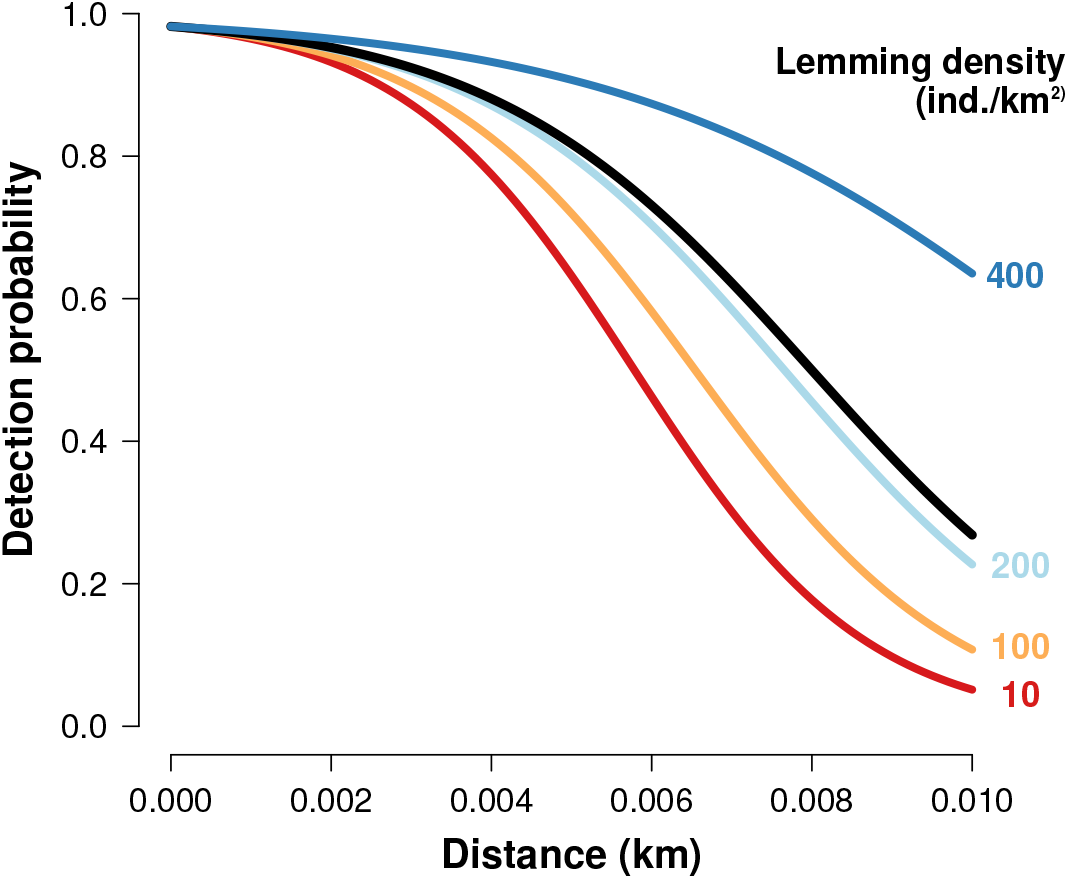
Probability that a lemming is detected by an arctic fox along a gradient of lemming density. The black line represents the average detection probability. The color gradient indicates the range of detection functions used to explore density dependence in detection probability.

